# Anti-CRISPR proteins trigger a burst of CRISPR-Cas9 expression that enhances phage defense

**DOI:** 10.1101/2023.10.22.562561

**Authors:** Rachael E. Workman, Marie J. Stoltzfus, Nicholas C. Keith, Chad W. Euler, Joseph Bondy-Denomy, Joshua W. Modell

**Affiliations:** Department of Molecular Biology & Genetics, Johns Hopkins University School of Medicine, Baltimore, MD 21205, USA; Department of Medical Laboratory Sciences, Hunter College, CUNY, New York, New York, USA; Department of Microbiology and Immunology, Weill Cornell Medical College, New York, New York, USA; Department of Microbiology and Immunology, University of California, San Francisco, San Francisco, CA 94158, USA; Quantitative Biosciences Institute, University of California, San Francisco, San Francisco, CA 94158, USA; Innovative Genomics Institute, Berkeley, CA, USA

## Abstract

CRISPR-Cas immune systems provide bacteria with adaptive immunity against bacteriophages, but they are often transcriptionally downregulated to mitigate autoimmunity. In some cases, CRISPR-Cas expression increases in response to a phage infection, but the mechanisms of induction are largely unknown, and it is unclear whether induction occurs strongly and quickly enough to benefit the bacterial host. In *S. pyogenes*, Cas9 is both an immune effector and autorepressor of CRISPR-Cas expression. Here, we show that phage-encoded anti-CRISPR proteins relieve Cas9 autorepression and trigger a rapid increase in CRISPR-Cas levels during a single phage infective cycle. As a result, fewer cells succumb to lysis leading to a striking survival benefit after multiple rounds of infection. CRISPR-Cas induction also reduces lysogeny, thereby limiting a route for horizontal gene transfer. Altogether, we show that Cas9 is not only a CRISPR-Cas effector and repressor, but also a phage sensor that can mount an anti-anti-CRISPR transcriptional response.

## Introduction

CRISPR-Cas loci encode adaptive immune systems that provide prokaryotic hosts with protection against foreign genetic elements, including bacteriophages (phages). CRISPR-Cas systems acquire short DNA-based memories from phages and store them in the CRISPR array as “spacers” to establish a record of past infection events^1^ (Fig. 1A). Spacers are transcribed into *crRNA* guides which direct Cas effectors to detect and disrupt complementary phage nucleic acid targets during a subsequent infection in a process called “interference”. Although CRISPR immunity provides protection against phages, there is a growing appreciation that CRISPR systems can be costly to maintain^2–7^. To mitigate these costs, CRISPR-Cas expression is often repressed. In some bacteria, transcription factors outside the CRISPR-Cas locus can regulate CRISPR-Cas expression in response to changes in temperature, quorum-sensing, growth phase, and metabolism^8–12^. As these conditions may or may not portend a phage infection, an open question is whether CRISPR-Cas systems can be directly regulated by phage themselves. There are several reports that the abundance of CRISPR-Cas components increases during a phage infection^13–16^, but in most cases the regulatory mechanisms are unclear, and it is unknown if the observed changes can take place quickly enough to affect immunity.

**Figure 1.**
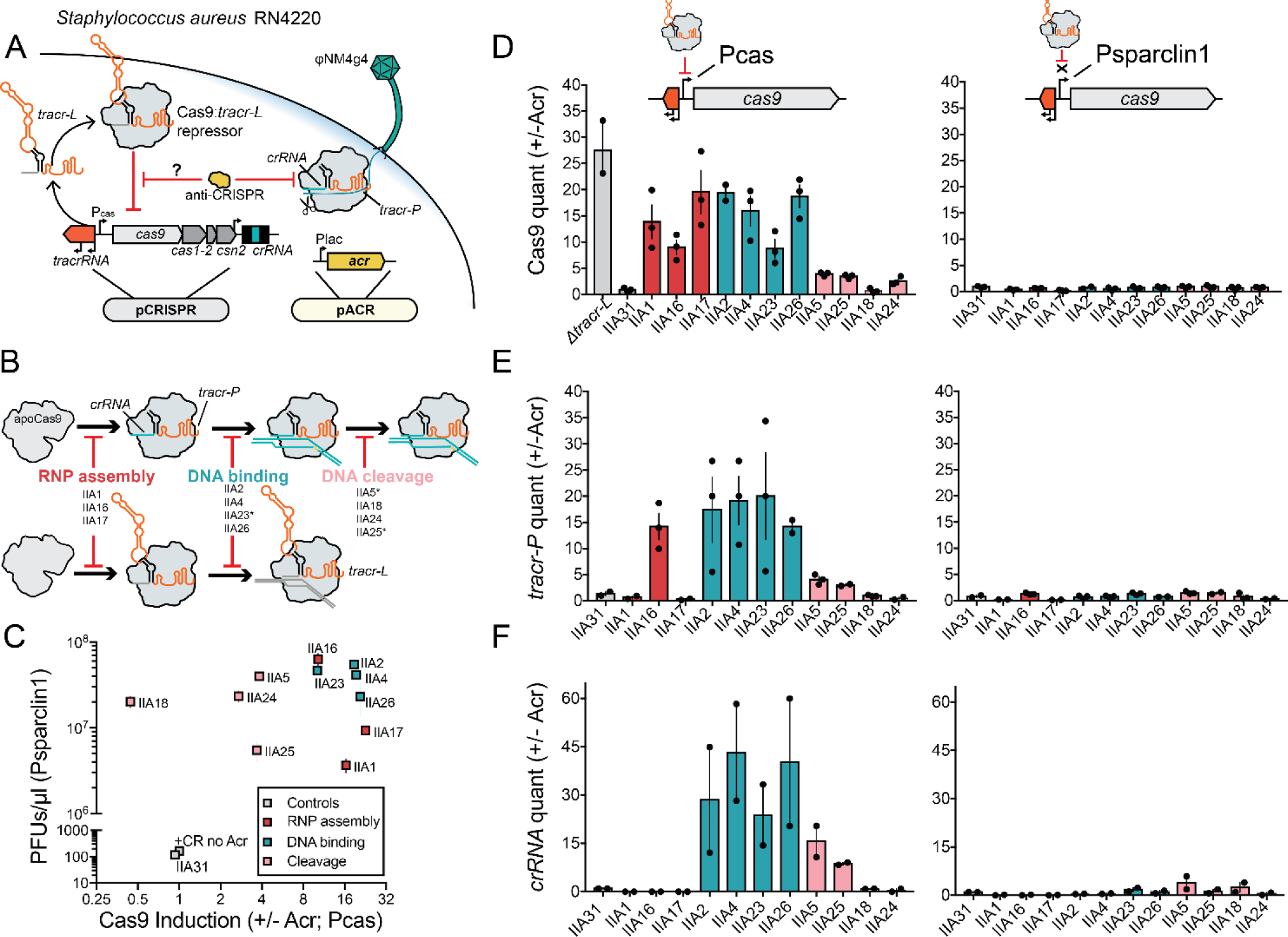
Anti-CRISPRs induce CRISPR-Cas expression through disruption of the Cas9:*tracr-L* repressor. A) Schematic of the model system in which *S. aureus* RN4220 hosts a plasmid with the complete type II-A CRISPR-Cas system from *S. pyogenes* (pCRISPR) and a second plasmid with an IPTG inducible anti-CRISPR (pACR). B) The indicated anti-CRISPRs inhibit Cas9 interference through RNP assembly, Cas9 DNA binding, Cas9 cleavage, or a combination of cleavage and binding inhibition (*). C) X-axis, Cas9 levels were measured by Western blot in Pcas cells following a 120 min IPTG induction and normalized to a no Acr control. Y-axis, PFUs/μl were measured in a top agar interference assay with φNM4γ4 in Psparclin1 cells grown in IPTG to express the indicated Acr. D) Cas9 Western blot quantification of Pcas or Psparclin1 cells following a 120 min IPTG induction for the indicated Acrs, normalized to a no Acr control. E-F) Quantification of Northern blots probing for *tracr-P* and *crRNA* in cells treated as in (C), normalized to a no Acr control. All experiments throughout are performed in biological duplicate or triplicate, error bars are standard error (*p < 0.05).

In type II-A CRISPR-Cas systems, the Cas9 effector binds to a dual-RNA guide consisting of a *crRNA* and a transactivating *crRNA* (*tracrRNA*) scaffold. The CRISPR array is transcribed as a single precursor *crRNA* (*pre-crRNA*) which contains spacers surrounded by repeating sequences^17^ (Fig. 1A). *tracrRNAs* base-pair with *pre-crRNA* repeats and the two RNAs are together cleaved by RNAseIII, producing individual *crRNA*s and a processed *tracrRNA* (*tracr-P*). In *S. pyogenes*, *tracrRNA* is transcribed from two promoters, leading to the production of a short-form and long-form (*tracr-S* and *tracr-L*, respectively), which can both base-pair with a *pre-crRNA* to perform processing and interference^7,17,18^. We found that *tracr-L* can also fold into a single-guide RNA that directs Cas9 to target and repress the Cas gene promoter (Pcas), leading to a reduction in *cas* gene, *tracrRNA*, and *crRNA* expression^18^. As a result, *S. pyogenes* cells express low levels of Cas9 RNP which reduces auto-immunity in the absence of phage but also limits immunity in the presence of phage^18^. We therefore wondered if physiological cues could trigger de-repression of *tracr-L* and provide hosts with a mechanism to increase CRISPR-Cas expression as needed, such as in response to phage infection. We hypothesized that inhibition of the Cas9:*tracr-L* repressor complex could be accomplished by any factor that more generally inhibits Cas9 function, leading us to investigate phage-encoded anti-CRISPR proteins^19^.

Anti-CRISPRs (Acrs) are small proteins, typically encoded on phage or mobile genetic elements, that inhibit CRISPR-Cas immunity^20^. Phage-encoded Acrs are expressed from a strong promoter early during an infection and are later repressed by a co-operonic anti-CRISPR associated gene (Aca), which prevents the deleterious overexpression of downstream genes^21–23^. Acrs of type II CRISPR-Cas9 systems can affect (i) formation or stability of the Cas9:*tracr-P:crRNA* ribonucleoprotein (RNP) complex ^24–26^, (ii) binding of Cas9 RNP to target DNA^27,28^, or (iii) Cas9 dsDNA cleavage activity^26,27^. It was recently shown that heterologous Acr expression from a plasmid could alleviate CRISPR-Cas auto-repression in type I and type V systems^29^, but it is unclear whether this occurs in type II systems and critically, whether CRISPR-Cas induction occurs strongly and quickly enough to benefit cells during a phage infection.

Here we establish that anti-CRISPRs with diverse mechanisms can de-repress the Cas operon through disruption of the Cas9:*tracr-L* repressor. We perform an in-depth characterization of CRISPR-Cas components during Acr expression and uncover new mechanistic insights for several Acrs. Next, we show that Acr-expressing phages (Acr-phages) can trigger a burst of CRISPR-Cas expression on short timescales that coincide with a single lytic cycle. This burst of expression protects *Streptococcus pyogenes* from phages by (i) limiting the number of infected cells which succumb to lysis or lysogeny and (ii) enhancing the survival of re-infected cells. Our work demonstrates a novel and direct mechanistic link between phage infection and CRISPR-Cas expression and highlights a strategy hosts can employ to maximize CRISPR-Cas immunity while minimizing auto-immune costs. As such, the *tracr-L* auto- regulatory circuit represents a weapon in the bacteria-phage arms race that serves as a countermeasure for CRISPR-Cas hosts against the emergence of phage-encoded Acrs.

## Results

### Anti-CRISPRs induce Cas expression through inhibition of the Cas9*:tracr-L* repressor

We selected a panel of 12 Acrs found in *Streptococcal* or *Listerial* phages that are known to inhibit *Streptococcus pyogenes* Cas9 (SpyCas9) by diverse mechanisms (Supplemental Data 1), including impacting Cas9 RNP formation through Cas9 stability or synthesis (AcrIIA1)^25^ or RNP assembly (AcrIIA16, AcrIIA17)^24,26^, target DNA binding (AcrIIA2, AcrIIA4, AcrIIA5, AcrIIA25, AcrIIA26)^27,28,30^, and target DNA cleavage (AcrIIA5, AcrIIA25, AcrIIA18, AcrIIA24)^26,27,30^ (Fig. 1B). We also included AcrIIA23 which inhibits SpyCas9 through an unknown mechanism^22^ and AcrIIA31, which inhibits *Streptococcus thermophilus* Cas9 (StCas9) but not SpyCas9^27^.

To study the impact of Acrs on CRISPR-Cas expression, we used a heterologous system in which *S. aureus* cells harbor one plasmid expressing the *S. pyogenes* CRISPR-Cas system and a second plasmid expressing Acrs from an IPTG-inducible promoter (Fig. 1A). To confirm Acr function and measure relative Acr strengths, we introduced a spacer targeting phage ΦNM4g4 into the CRISPR array and performed a phage interference assay (Fig. 1C). To eliminate the confounding influence of CRISPR-Cas induction on interference, we replaced the Cas operon promoter Pcas with a constitutive promoter Psparclin1 (Supplemental Data 1) that is not repressed by *tracr-L*. Each Acr inhibited Cas9 interference by between 4-6 orders of magnitude, except for the negative control AcrIIA31 which did not affect interference.

We next asked whether Acrs impacted expression of the *S. pyogenes* CRISPR-Cas system. Cas9 protein levels were quantified by Western blot following 2 hours of Acr induction. We observed a striking 10-20- fold increase in Cas9 levels upon expression of Acrs that inhibit RNP formation and target DNA binding (Fig. 1D, S1A). An intermediate 2-5-fold increase in Cas9 levels was observed for Acrs that inhibit Cas9 cleavage, except for AcrIIA18, which did not significantly affect Cas9 expression. While the DNA binding inhibitors were all strong Cas9 inducers and strong interference inhibitors, the RNP and cleavage inhibitors did not show a strong correlation between induction and inhibitory strength (Fig. 1C), suggesting that the degree of induction may depend on mechanistic details that distinguish Acrs within each class.

Next, we sought to understand whether Acr-dependent CRISPR-Cas induction relies on displacement of the Cas9:*tracr-L* repressor from Pcas. When the CRISPR-Cas system was expressed from the constitutive promoter Psparclin1, Cas9 levels did not increase following Acr induction for any inducing Acr (Fig. 1D, S1B), suggesting that Acrs affect Cas9 expression at the level of transcription. Cas9 levels decreased for AcrIIA1 in Psparclin1 cells (Fig. S1B), consistent with its known role in inhibiting Cas9 synthesis or accelerating its degradation^25,31^. This decrease was likely masked by Cas9 induction in Pcas cells.

Unexpectedly, Cas9 levels also decreased for AcrIIA17 in Psparclin1 cells, suggesting that it is a bifunctional inhibitor that affects RNP assembly as well as Cas9 synthesis and/or degradation. To confirm that Cas9-inducing Acrs affect the activity of the Pcas promoter, we constructed a plasmid expressing Cas9, *tracr-L*, and a Pcas-GFP transcriptional reporter. Following Acr induction, we observed increases in Pcas activity proportional to the increases in Cas9 protein levels for most Acrs (Fig. S1C).

Taken together, these observations indicate that Acrs can regulate CRISPR-Cas expression by inhibiting the Cas9:*tracr-L* repressor and enhancing transcription of the Cas gene operon.

Acrs are typically studied in the context of Cas9 bound to an *sgRNA*. To gain further mechanistic insights into how Acrs can affect native CRISPR-Cas RNAs, we monitored *crRNA* and *tracrRNA* levels following Acr induction in both Pcas and Psparclin1 cells. We performed Northern blots using a probe that hybridizes to the 3’ end of *tracrRNA* and recognizes *tracr-P*, *tracr-L* and *tracr-S*, and a probe that hybridizes to the *crRNA* repeat and recognizes *pre-crRNAs* and *crRNAs* (Fig. S1A-B). When the DNA-binding inhibiting Acrs were expressed, *tracr-P* and *crRNA* levels increased, mirroring those of Cas9, likely because Cas9 stabilizes these RNAs^18^. AcrIIA23 largely phenocopied the DNA-binding inhibitors suggesting that it is a member of this class. For the RNP formation inhibitors AcrIIA1 and AcrIIA17, *tracrRNA* and *crRNA* levels decreased (Fig. 1E-F, S1A) suggesting that Acr overproduction was sufficient to prevent pre-existing and newly synthesized Cas9 from binding to RNAs. The RNP inhibitor AcrIIA16 also led to a decrease in *crRNAs*, consistent with its known role in *sgRNA* degradation^24^. *tracr-P* unexpectedly increased by 10- fold, but several bands of unexpected sizes were observed on both the *tracrRNA* and *crRNA* Northern blots, casting doubt on the precise identity of the *tracr-P* band and suggesting that AcrIIA16 may cause degradation of both guide RNAs in the native system. AcrIIA5 and AcrIIA23 led to the preferential accumulation of the 39nt and 42nt processed *crRNAs* respectively, and both Acrs caused increases in *tracr-P* (Fig. 1E-F, S1A-B), suggesting that these Acrs could affect *pre-crRNA* processing. Collectively, we show that measuring Cas9 induction and native RNA expression levels can help classify new Acrs and provide novel mechanistic insights for previously characterized Acrs.

### CRISPR-Cas induction promotes bacterial survival during an Acr-phage infection

The above results indicate that Acr overproduction from a plasmid can trigger *S. pyogenes* Cas9 induction in a heterologous *S. aureus* host. Next, we asked if phage-encoded Acrs expressed from a native promoter could trigger Cas9 induction in time to benefit *S. pyogenes* cells during a phage infection. We selected phage A1, the only phage that has been experimentally validated in *S. pyogenes* CRISPR-Cas assays^32^. Because phage A1 does not host any identifiable Acrs, we engineered Acrs into a location where they are typically found within *Streptococcal* phage genomes (Fig. S2A-B). Acrs were expressed from the AcrIIA3 promoter (Pacr) found in the phiSTAB1 prophage within *S. pyogenes* strain STAB13021. This Acr locus contained a candidate Aca with 43.5% identity to aca11^27^, which we verified controlled expression of Pacr (Fig. S2C). Selected Acrs from the panel above were expressed from Pacr just upstream from the validated Aca to recapitulate the operonic and regulatory structure that is highly conserved across Acr-expressing phages. In this way, we introduced 9 Acrs into phage A1: 2 RNP inhibitors (AcrIIA16, AcrIIA17), 4 DNA binding inhibitors (AcrIIA2, AcrIIA4, AcrIIA23, AcrIIA26), and 3 cleavage inhibitors (AcrIIA5, AcrIIA18, AcrIIA25).

To quantify Cas9 induction during an Acr-phage infection, we infected *S. pyogenes* strain C13, a derivative of SF370 that lacks prophages^33^, with the Acr-expressing A1 variants (Fig. 2A). After 75 minutes of infection, the shortest time required for phage A1 to complete its lytic cycle in the absence of CRISPR targeting (Fig. S2D), we found that A1 expressing the inducing AcrIIA26 (A1-IIA26) led to a dramatic 6-7-fold increase in Cas9 levels, while cells infected with wild-type A1 or A1 expressing the non-inducing AcrIIA18 showed no change (Fig. 2B). The Cas9 induction patterns for the remaining *S. pyogenes* A1-Acr infections were generally consistent with data obtained from the *S. aureus* heterologous system (Fig. 1D, 2C), except for AcrIIA17, which induced Cas9 in the heterologous system but not in the native system. This discrepancy could be explained by lower levels of Cas9 induction and higher rates of degradation in the native system. We next assayed Cas9 levels every 15 minutes following infection and found that for some inducing Acrs, Cas9 levels rise quickly, as early as 15-30 minutes after the start of infection (Fig. S2E). Overall, these results suggest that phage-encoded Acrs can strongly and quickly trigger Cas9 induction in a time frame that could benefit cells during a phage infection.

**Figure 2.**
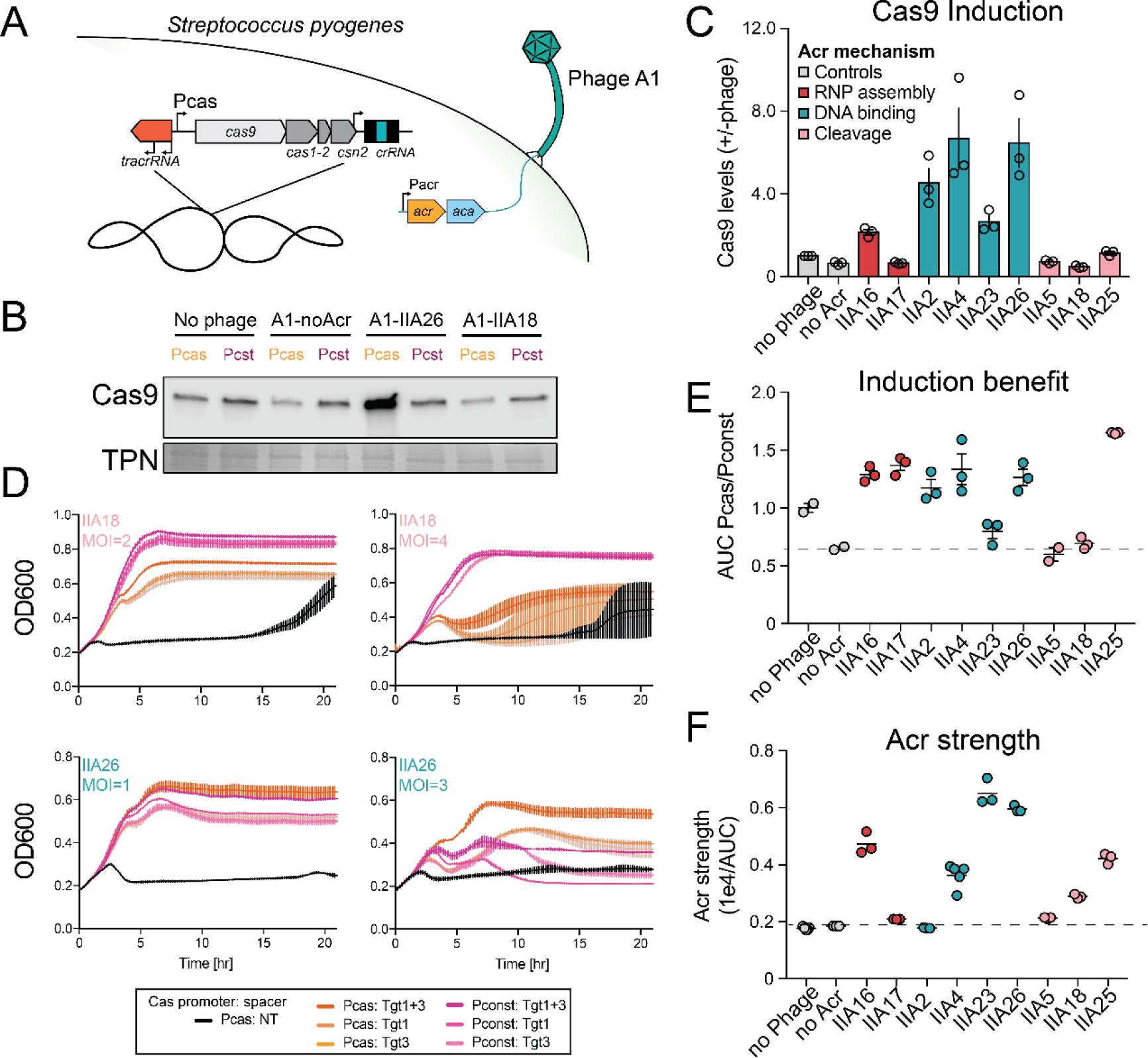
CRISPR-Cas induction during Acr-phage infection enhances bacterial survival. A) Schematic of the model system in which *S. pyogenes* strain C13 with spacers targeting phage A1 is infected with A1-Acr variants. B) Cas9 Western blot from Pcas and Pconst (Pcst) cells infected with WT A1 without an Acr, A1-IIA26, or A1-IIA18 at MOI=2 for 75 minutes. C) Quantification of a Cas9 Western blot with Pcas cells infected at MOI=2 for 75 minutes, normalized to an uninfected control strain. D) Growth curves of Pcas and Pconst cells infected with A1-IIA18 or A1-IIA26 at the indicated MOIs. E) Benefit of Pcas induction was quantified by dividing the area under the curve (AUC) of a 20 hour Pcas infection with that of a Pconst infection at the highest MOI allowing for survival of Pcas (MOIs provided in Supplemental Data 1). Values above the dotted line represent a benefit to Pcas induction. F) The strength of each Acr was quantified by calculating 1e4 divided by the AUC of a 20-hour infection of Pconst cells at MOI=4 with the corresponding Acr-phage.

To ask if Cas9 induction enhances *S. pyogenes* survival during an Acr-phage infection, we generated an uninducible control strain that expresses the Cas genes (i) from a promoter not regulated by *tracr-L* and (ii) at levels that approximate the uninduced state of wild-type cells. After many iterations of promoter testing and mutagenesis, we chose a promoter (Pconst) for which Cas9 levels were roughly 2-fold higher than WT in the absence of phage and were unchanged following a 75-minute infection with the strongly- inducing phage A1-IIA26 (Fig. 2B, S2F). Next, we generated six interfering *S. pyogenes* strains expressing Cas genes from Pcas or Pconst with either one (Tgt1, Tgt3) or two (Tgt1+3) spacers targeting phage A1. We infected each strain with Acr-expressing A1 variants and measured cell growth every 10 minutes for 24-hours. Pconst cells survive the non-inducing phage A1-IIA18 better than Pcas, likely because of the aforementioned 2-fold increase in basal CRISPR-Cas expression (Fig. 2D). Despite this Pconst advantage, Pcas cells survive better than Pconst when challenged with the inducing phage A1-IIA26 (Fig. 2D), suggesting that Cas9 induction enhances CRISPR-Cas defense. We quantified an “induction benefit” metric for the remaining Acr-phages by dividing the area under the curve for Pcas infections by that of a Pconst infection under identical infection conditions and found a significant benefit for the strong inducers (AcrIIA2, AcrIIA4, and AcrIIA26) (Fig. 2E, S2G). A strong induction benefit was also observed for AcrIIA17, for which degradation in the absence of induction leads to lower Cas9 levels in Pconst cells (Fig. 2E, S2F). Unexpectedly, a strong benefit was also observed for the mild-inducers IIA25 and IIA16, suggesting mechanistic differences that require further study.

Lastly, we asked whether the inhibitory strength of a given Acr dictates the intensity of Cas induction in the native host. To quantify Acr strength, we generated 24-hour growth curves of Pconst cells infected by each Acr-phage at MOI=4 and calculated area under the curve (Fig. 2F). As in the heterologous system, we found that each Acr mechanistic class included strong and weak interference inhibitors, and that Pcas benefit correlated well with induction potential and poorly with Acr strength (Fig. 2C,E-F).

### The benefits of CRISPR-Cas induction emerge after multiple lytic infective cycles

Acr-driven CRISPR-Cas induction benefits bacterial cells during a 24 hour infection (Fig. 2D-E), but when during the infection are the benefits of induction realized? Previous research has shown that Acr-phages often must complete multiple infections to successfully replicate and lyse a CRISPR-Cas host with a spacer targeting that phage^34,35^. In this scenario, pre-existing Cas9 RNPs can clear a single infecting phage; however, the Acrs synthesized during this failed infection immunocompromise the host such that an invading phage can successfully lyse the cell during a subsequent or second simultaneous infection.

We therefore wondered whether CRISPR-Cas induction can increase survival against an Acr-phage i) during a single infection, ii) during a simultaneous infection (i.e. MOI > 1), and/or iii) during multiple rounds of infection.

We first asked whether CRISPR-Cas induction could bolster bacterial survival during an infection by a single Acr-phage. We infected Pcas and Pconst cells at MOI = 0.1 with either the strongly inducing A1- IIA26 phage or non-inducing A1-IIA18 phage and measured the number of cells that lysed in an infective centers assay. Regardless of the spacer content, a slightly higher proportion of Pconst cells survived relative to Pcas for A1-IIA18 (Fig. S3A,C), reflecting the higher basal levels of Cas9 in the Pconst strain.

Similar results were observed for AI-IIA26 (Fig. S3B-C), suggesting that Cas9 induction does not appreciably enhance bacterial survival during a single infection. We quantified infective centers again at MOI = 3, which allows for simultaneous infections but still measures a single temporal infectious cycle. Under these conditions, while Pconst cells maintained an advantage over Pcas during an A1-IIA18 infection, Pcas and Pconst cells survived similarly during an A1-IIA26 infection (Fig. 3A), suggesting that Cas9 induction enhances survival during simultaneous infections with an inducing phage. We next measured the number of phages generated during a single round of infection at MOI = 3. Consistent with the cell survival assay, phage survival was enhanced in Pcas relative to Pconst for Al-IIA18 but was roughly equal for Al-IIA26 (Fig. 3B). We calculated the number of phages released per lysing cell (burst size) and found that it was the same for Pcas and Pconst during both A1-IIA18 and A1-IIA26 infections (Fig. S3D). This suggests that under these conditions, CRISPR-Cas induction affects the number of cells that burst more than the number of phages released by each burst.

**Figure 3.**
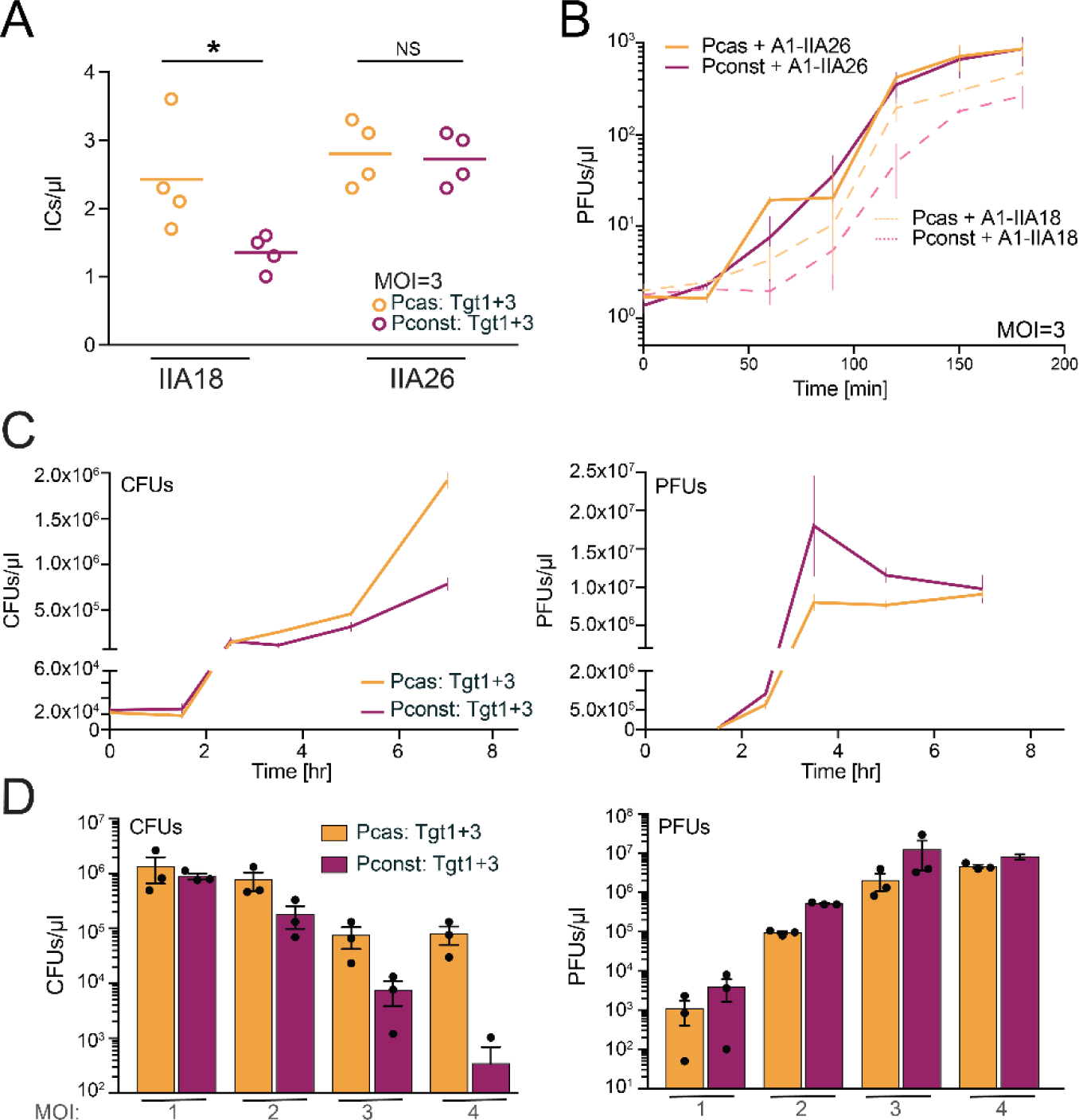
The benefits of CRISPR-Cas induction emerge upon multiple phage infective cycles. A) Infective centers assay, Pcas and Pconst cells were infected with A1-IIA18 or A1-IIA26 at MOI=3 and the number of infective centers per μl was calculated. B) One-step growth curve (OSGC) assay for Pcas and Pconst cells infected at MOI=3 with phage A1-IIA26 or A1-IIA18. C) Quantification of plaque forming units (PFUs) and colony forming units (CFUs) for Pcas or Pconst cells after an infection with phage A1-IIA26 at MOI = 3 at the indicated time points. D) PFUs and CFUs were calculated 24 hours after infection of Pcas or Pconst cells with A1-IIA26 at the indicated MOIs.

To test the benefit of induction after multiple consecutive rounds of infection, we infected Pcas and Pconst cells with the inducing phage A1-IIA26 at MOI=3, then plated for PFUs and CFUs over time. When a single lytic cycle had completed at 2.5 hours, cell survival and phage replication were roughly equal (Fig. 3C) despite the pre-induced advantage of Pconst over Pcas cells. CFUs increased in both cultures by roughly 7-fold from t=1.5h to t=2.5h, roughly coinciding with the completion of the first lytic cycle, likely because phage lysis can cause de-chaining of *Streptococcal* cells^36^. By 3.5 hours, during the second round of infections, induced cultures showed a slightly higher number of CFUs but dramatically fewer PFUs. By 7 hours, likely during the third wave of infections, a significant survival benefit emerged for induced cells, although the number of PFUs had equilibrated between the two strains (Fig.3C). These data demonstrate that under these conditions, (i) a reduction in phage replication precedes a significant boost in cell survival and (ii) the benefits of Cas9 induction are compounded after multiple infective rounds.

Our data show that the benefit of Cas9 induction may be MOI-dependent, as cell survival is enhanced after a single lytic cycle at MOI=3 (Fig. 3A-B) but not at MOI=0.1 (Fig. S3A-C). To explore whether the benefit of Cas9 induction is generalizable across infection conditions, we performed a 24-hour infection of Pcas and Pconst cells using phage A1-IIA26 at multiple MOIs (Fig. S3E) and quantified endpoint PFUs and CFUs (Fig. 3D). We found that as MOI increased, the benefit of induction for cell survival increased as Pcas cells outperformed Pconst cells. Curiously, although cell survival was similar at low MOIs, induction did reduce the number of phages present at the end of the timecourse (Fig. 3D). These data indicate that induction benefits CRISPR-Cas populations during lytic infections, by increasing cell survival and/or limiting phage replication across all MOIs tested.

### CRISPR-Cas induction reduces lysis and lysogeny by a native, temperate Acr phage

All known Acr-containing *S. pyogenes* phages are temperate, meaning they can either lyse the infected host or integrate into the host genome as a dormant prophage during “lysogeny”. Although phage A1 is reported to be temperate^32^, we were unable to obtain A1 lysogens and as mentioned, A1 does not contain Acrs.

To ask whether Cas9 induction impacted lysis and lysogeny of a native Acr-phage, we used *S. pyogenes* phage AP1.1-spec^22^, which natively encodes AcrIIA23 and contains an engineered spectinomycin-resistance gene to enable selection for lysogens (Fig. S4A). We programmed Pconst or Pcas *S. pyogenes* cells with a single spacer targeting AP1.1 (Sp1), and we generated AP1.1-spec variants in which AcrIIA23 was deleted or replaced with the non-inducing AcrIIA18 or strongly inducing AcrIIA26. While we were unable to achieve sufficient AP1.1 titers to lyse CRISPR-targeting cells in liquid culture, we were able to obtain PFUs in a top-agar interference assay. As expected, the non-inducing phage AP1.1-IIA18 replicated more efficiently on Pcas top agar lawns than Pconst, consistent with the higher basal Cas9 expression in Pconst cells (Fig. 2B). However, the advantage for Pconst cells was significantly diminished when AcrIIA18 was replaced with AcrIIA23 or AcrIIA26, both capable of inducing Cas expression (Fig. 4A, S4B). This suggests that the lytic cycle of the native temperate phage AP1.1 is sensitive to CRISPR-Cas induction.

**Figure 4.**
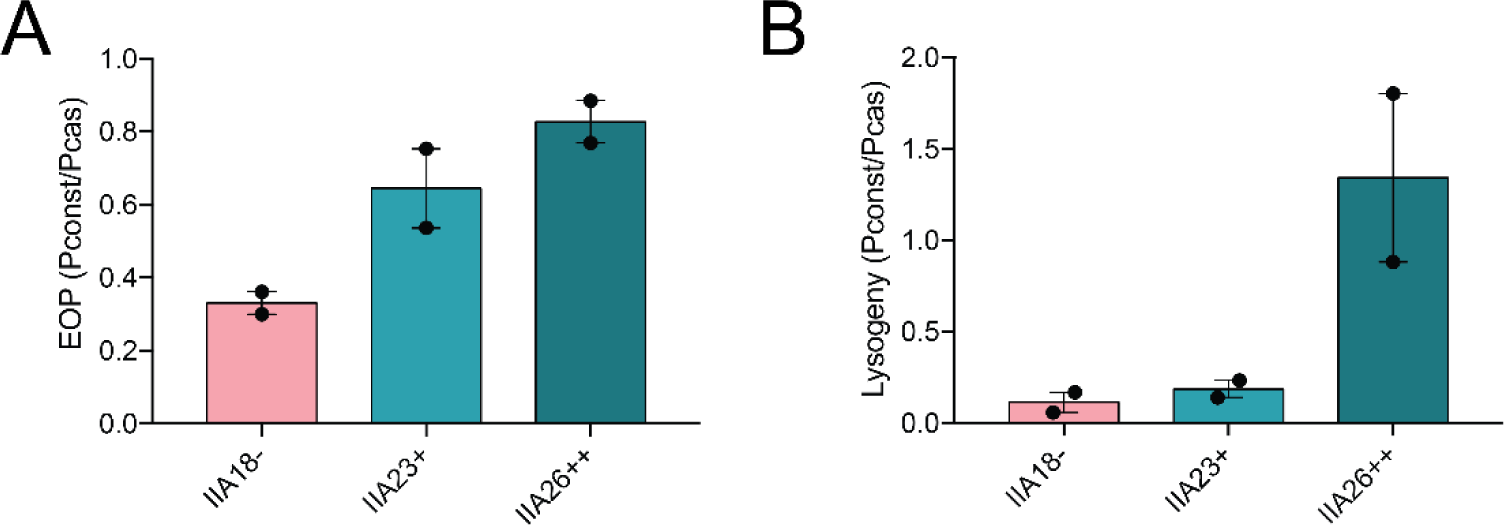
CRISPR-Cas induction inhibits AP1.1 lysis and lysogeny. The ratio of efficiency of plating (A) and lysogeny rates (B) for Pconst cells divided by that of Pcas cells (both Sp1) infected with the indicated AP1-Acr variants (-, non-inducer; +, mild inducer; ++, strong inducer). Individual EOP and lysogeny values for Pcas and Pconst are provided in Fig. S4.

We next assayed the potential of the AP1.1 phages to form lysogens by infecting cells at a low MOI (∼0.0001) and plating on media supplemented with spectinomycin. Significantly more lysogens were observed for Pcas relative to Pconst for both the non-inducing phage AP1.1-IIA18 and wild-type phage AP1.1-IIA23, again reflecting the higher basal Cas9 expression in Pconst cells. In contrast, lysogen formation of the strongly inducing phage AP1.1-IIA26 was roughly equivalent in Pcas and Pconst cells (Fig. 4B, S4C). This suggests that CRISPR-Cas induction can limit lysogeny of Acr-expressing phages during a single infective round with a single phage.

Altogether, our data support a model (summarized in Fig. 5) whereby inducing Acr-phage infections can quickly lead to de-repression of the Cas operon, resulting in reduced lysogeny and cell lysis within a single infective round. The benefits of limited lytic replication and increased Cas9 expression are then compounded after successive rounds of infection.

**Figure 5.**
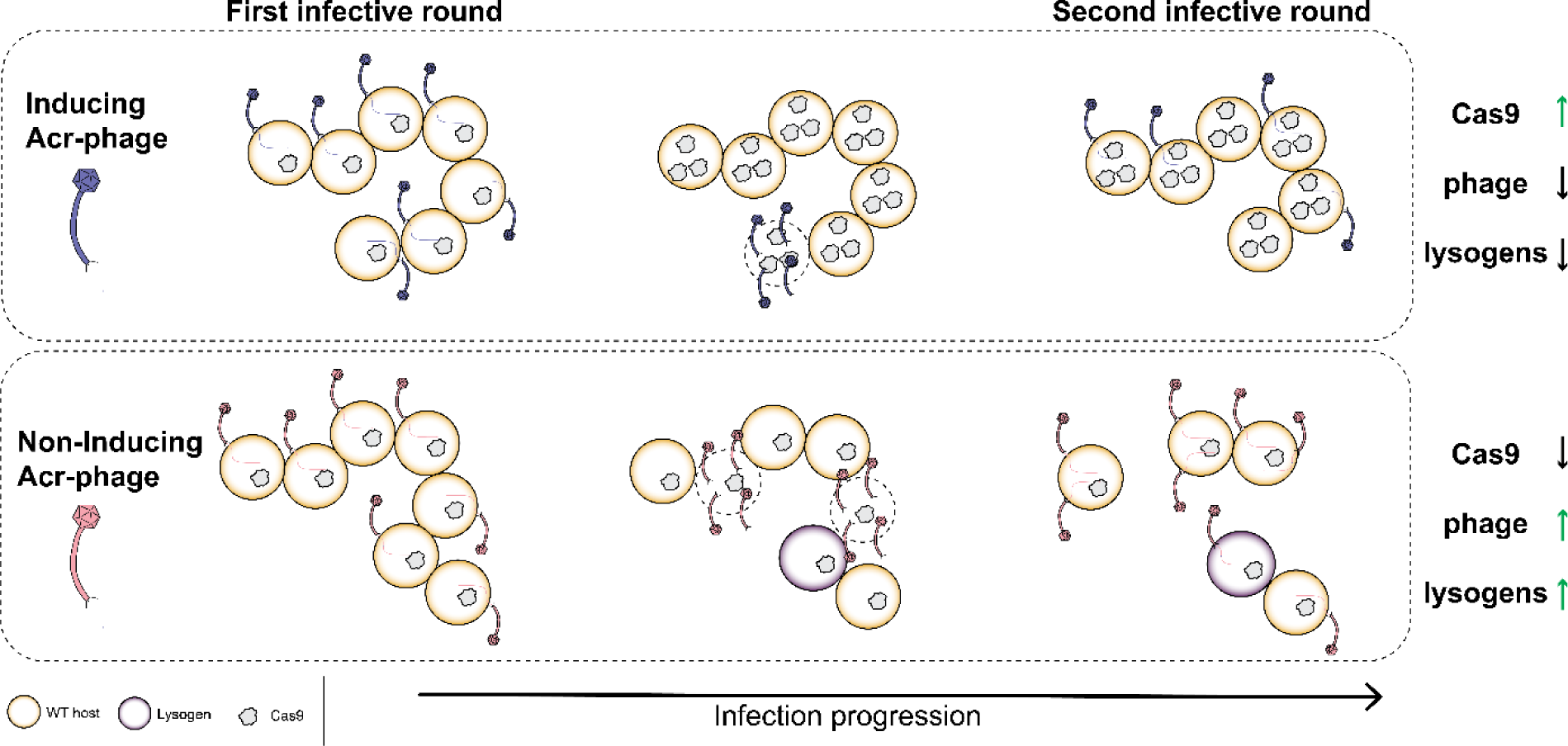
Cas9 defends against bacteriophage anti-CRISPRs by transiently inducing CRISPR-Cas expression. Our data support a model in which CRISPR-Cas9 systems can respond to an Acr-phage infection by quickly increasing CRISPR-Cas expression, resulting in reduced lytic replication and lysogenic conversion within a single infective round. The benefits of limited lytic replication and increased Cas9 expression are compounded after successive rounds of infection, resulting in a considerable cell survival benefit.

## Discussion

### CRISPR-Cas systems are dynamically regulated

Because CRISPR-Cas overexpression leads to fitness defects for the bacterial host^6,3,18^, many CRISPR-Cas systems are repressed by intrinsic or extrinsic factors^8–12^. Whether and how CRISPR-Cas repression can be relieved as needed, for example during a phage infection, is an open question. In *Streptococcus pyogenes,* the regulatory RNA *tracr-L* reprograms Cas9 from an immune effector into a transcriptional repressor of the Cas operon promoter Pcas, dampening immunity while also reducing fitness costs of CRISPR-Cas overexpression. Here, we report that phage-encoded anti-CRISPRs can transiently trigger CRISPR-Cas overexpression by relieving Cas9:*tracr-L* repression. Using both heterologous and native systems, we demonstrate that Acrs from all known mechanistic classes can trigger Cas9 induction, and the intensity varies depending on the mechanism of the Acr (Fig.1).

Our work is the first to provide a mechanistic understanding of phage-responsive CRISPR-Cas activation for the Cas9-encoding, type II CRISPR-Cas systems. We find that *tracr-L* is not solely a transcriptional repressor, but also a responsive sensor element and anti-phage countermeasure. Interestingly, it is not the only Cas regulator which moonlights as a responsive sensor element. In the archaeal *Sulfolobus islandicus* type I-A system, the interference complex Cascade interacts with a CRISPR-Cas transcriptional repressor Csa3b. During a phage infection, sequestration of Cascade to the phage target leads to de-repression of Csa3b^37^. Additionally, during the preparation of this manuscript, a study of several type I-B, I-E, and V-A systems found mini-arrays encoding *cas*-regulating RNAs (CreRs) that can be relieved by the heterologous expression of Acrs^29^. CRISPR-Cas regulation by phage-encoded Acrs may therefore be a general property of CRISPR-Cas systems with auto-regulatory circuits. Curiously, increases in Cas9 levels following phage infection have been reported in several species which do not encode *tracr-L* or other obvious auto-regulatory RNAs^13–15^. It remains unknown whether these systems possess alternative means of autoregulation or extrinsic phage-responsive regulators.

### Acr-dependent Cas induction is rapid and beneficial to hosts

A crucial unanswered question is whether CRISPR-Cas induction during a phage-infection occurs quickly and strongly enough to impact host survival. In the context of an Acr-phage, can new Cas9 RNP synthesis outcompete Acr synthesis before the lytic cycle completes? *S. pyogenes* phage A1 begins to replicate exponentially within 15 minutes after infection, has a latency period of ∼60 minutes, and produces ∼20 virions per cycle^32^ in the absence of host interference (Fig. S2D). In contrast, the CRISPR-Cas operon must be de-repressed, transcribed, translated, *crRNAs* and *tracrRNAs* must be processed, and Cas9 RNPs must find and cleave their targets. Remarkably, we show that Cas induction through *tracr-L* de-repression is rapid, as Cas9 levels can double in as little as 15 minutes after the start of phage infection and increase as much as 10-fold within 75 minutes.

To quantify the benefits of CRISPR-Cas induction, we sought to express the CRISPR-Cas system from a *tracr-L* independent promoter at levels similar to the un-induced native system. Ultimately, after much trial and error, we settled on the Pconst promoter which expressed Cas9 at levels 2X higher than uninduced wild-type cells (Fig. 2B). Practically, this means that in assays comparing Pcas and Pconst cells, Pcas cells are at an initial disadvantage which is maintained during exposure to phages with non-inducing Acrs (Fig. 2E). In contrast, when Pcas and Pconst cells break even against a phage with an inducing Acr (Fig. 3A, S4B-C), we interpret this as Pcas “winning” relative to Pconst and attribute that advantage to Cas induction.

We show that a small cellular benefit of induction in a single phage lytic cycle can nonetheless lead to a significant disparity in phage replication, resulting in (i) lower MOIs and (ii) higher levels of Cas9 RNP during subsequent infections. As a result, the benefits of induction are most pronounced during simultaneous infections (Fig. 3D, S3E) and over longer periods of time (Fig. 2D-E, 3C). This strategy of preparing cells for future infections dovetails nicely with previous work showing that phage-encoded Acrs often exert their effects through cooperation^34,35^. In this model, a single Acr-phage is killed by the CRISPR-Cas system, but not before depositing Acrs that leave host cells immunocompromised and more easily overcome by subsequent Acr-phage infections. By triggering CRISPR-Cas induction, *tracr-L* therefore protects against future infections by attenuating the immunocompromised state. During temperate AP1.1 phage infection, a benefit of induction can be realized within a single infection for both lytic replication and lysogenic conversion (Fig. 4B), in contrast to lytic phage A1 infections for which survival benefits were more noticeable at higher MOIs (Fig. 3D, S3E). The basis for this discrepancy between phage A1 and AP1.1 could be due to differences in the strength of the spacers studied, the timing and intensity of Acr expression, as well as other phage lifestyle differences.

### Evolutionary perspectives on *tracr-L* and Acr-phages

The antagonism of Acr-phages by *tracr-L* reveals another facet of the Red Queen hypothesis, in which bacteria and phage each continuously evolve defenses and counter-defenses in order to avoid extinction. We note that while *tracr-L* enhances CRISPR-Cas protection against phages with inducing Acrs, it does not render these Acrs obsolete. Rather, phages with inducing Acrs still perform dramatically better than phages lacking an Acr, and they are never extinguished from the population. Why then would phages carry an inducing Acr instead of a non-inducing Acr? Inducing Acrs tend to be among the most potent inhibitors of Cas9 interference (Fig. 2C), which could provide phages with an advantage during infections of the ∼57% of Streptococci that do not encode *tracr-L*^18^. Furthermore, nearly all Acrs tested provide some level of CRISPR-Cas induction (Fig. 1D, 2C, S1A), so there may be other evolutionary constraints that limit the utility of true, non-inducing Acrs like AcrIIA18.

To study the benefit of CRISPR-Cas induction, we cloned diverse Acrs into two experimental systems. First, we expressed Acrs within a native *Streptococcal* Acr regulon and inserted this locus into the corresponding region of phage A1. Second, we exchanged the native AcrIIA23 within the temperate phage AP1.1 with strongly- and weakly-inducing Acrs. In each case, we observed a benefit for CRISPR-Cas induction, suggesting that the principle likely applies more broadly. AcrIIA23 is itself a mild-inducer (Fig. 2C) that confers an even milder benefit of induction (Fig. 2E). IIA23 may therefore be an ideal Acr for AP1.1, allowing the phage to inhibit Cas9 interference while limiting CRISPR-Cas induction. While we were unable to obtain a native phage with an Acr that confers a strong advantage for CRISPR-Cas inducing cells, we note that AcrIIA16, AcrIIA17, and AcrIIA26 were found in prophages within *Streptococcal* species that can contain *tracr-L*^18^, which include *S. pyogenes*, *S. suis*, and *S. parasanguinus* (Supplemental data 1).

Quantifying the short and long-term “benefit” of Cas induction becomes more complicated when considering temperate phage infection. Lysogeny can be costly for bacteria due to the disruption of genes surrounding the integration site, the increased genomic size, and the threat of lysis during DNA damage ^38,39^. However, lysogeny is also a mode for the horizontal transfer of beneficial genes, like those involved in antibiotic-resistance, phage defense, and pathogenicity^40,41^. CRISPR-Cas induction may therefore provide short-term benefits by reducing lysis and reducing genome expansion while limiting the potential for adoption of new genetic material. Another question is how Acr expression from a prophage would affect CRISPR-Cas induction and function. Many Acrs are constitutively expressed from prophages, likely at higher levels than during a transient phage infection. To date, AP1.1 is the only prophage with a characterized and inducing Acr found within a genome that also contains *tracr-L,* and its Acr (AcrIIA23) is uncharacteristically not expressed in the lysogen^22^. Understanding the relationship between prophages and CRISPR-Cas induction will therefore require the characterization of additional *S. pyogenes* prophages and their Acrs.

### New insights into Acr mechanisms and fundamental CRISPR-Cas biology

Cas9 Acrs are actively studied both for their applications as gene editing regulators and for what they can teach us about Cas9. Acrs are typically tested in cleavage assays using Cas9:*sgRNA* and target binding assays using catalytically inactive Cas9 (dCas9). Here, we study Acr activity against the native CRISPR-Cas locus which offers several benefits. First, disruption of the Cas9:*tracr-L* repressor provides a measure of target DNA binding without the need for using engineered reporters or catalytically inactive Cas9 (dCas9), which can confound the interpretation of Acr mechanism^25^. Second, the native CRISPR-Cas locus enables the study of how Acrs affect *pre*-*crRNA* and *tracrRNA* processing, a step that is wholly absent when studying sgRNAs. Notably, while 39 and 42nt processed *crRNAs* are typically found in equivalent abundance (Fig. S1A-B), AcrIIA23 and AcrIIA5 expression leads to preferential accumulation of the 42nt or 39nt form, respectively. As yet, it is unclear how these two *crRNA* isoforms are naturally generated or whether they differ functionally. Similarly, *tracr-L*, *tracr-S*, and *tracr-P* are typically found in equal abundance, but AcrIIA1 and AcrIIA17 each lead to the specific depletion of *tracr-L* and *tracr-P*. Future studies will reveal whether these disparities are central to the inhibitory mechanism or side effects of an upstream activity. Third, we can uncover substantial mechanistic differences between inhibitors of the same “class”. For example, expression of all three RNP formation inhibitors (AcrIIA1, AcrIIA16, and AcrIIA17) lead to the loss of processed *crRNA*s, but AcrIIA16 expression is unique in that *tracrRNAs* and *pre-crRNAs* of intermediate lengths, possibly degradation products, increase in Pcas cells as Cas9 levels increase. This suggests that AcrIIA16 binding to Cas9 either does not exclude *tracrRNA* or *pre-crRNA* association, or that the Acr itself is sequestering these RNAs and preventing them from rapid turnover observed in the absence of Cas9 association.

Using the native CRISPR-Cas system, we assign AcrIIA23 to the class of target DNA binding inhibitors, although it shows several unusual phenotypes. As mentioned, it causes the preferential accumulation of the 42nt *crRNA*, suggesting that it influences *crRNA* binding or processing (Fig. S1A-B). Further, it is the weakest inducer in its class despite being the most potent DNA binding inhibitor as measured by interference against phage (Fig. 2E-F). More generally, we observed poor correlation between potency and induction across all Acrs (Fig. 2E-F), suggesting that Acrs may have different affinities for or activities against Cas9:*tracr-L* and Cas9:*tracr-P*:*crRNA*. We discovered that AcrIIA17 causes degradation of Cas9 (Fig. S1B, S2F) and that AcrIIA1 blocks RNP guide loading (Fig. S1A-B). AcrIIA5 is of interest as a broad-spectrum inhibitor of multiple Cas9 orthologs, but whether it blocks target DNA binding or cleavage has been a matter of debate. Our results support both mechanisms, and we speculate the discrepancies in earlier studies may be due to assay sensitivity and/or use of a heterologous host. We show that Acrs from all mechanistic groups can trigger a wave of CRISPR-Cas induction that benefits the bacterial cell.

Curiously, AcrIIA25 conferred the highest benefit of induction despite being a mild inducer (Fig. 2E-F). Whether this owes to the stoichiometry of the Acr with regard to Cas9 or other mechanistic details will require future studies.

Altogether, our data reveals the CRISPR-Cas autoregulator *tracr-L* represents a dimmer switch of CRISPR immunity that minimizes autoimmunity but also decreases phage defense. Here, we show that Acrs can flip that switch allowing bacteria to sense and respond to a phage infection. Future studies of natural *Streptococcal* communities would allow for a more nuanced study of how Acr-phages and CRISPR-Cas regulatory genes interact and coevolve.

## Acknowledgements

We would like to thank Adair Borges for early discussions during conceptualization of this study. We thank Vince Fischetti for reagents including the PlyC expression vector, Kevin McIver for allelic exchange vectors, and all the members of the Modell Lab for helpful feedback and suggestions. We thank David Mohr and the GRCF High Throughput Sequencing Center for assistance with NGS experiments. Funding was provided by a startup package from the Johns Hopkins School of Medicine, NIH NIGMS R35GM142731, and the Rita Allen Foundation 90094894.

## Author Contributions

R.E.W. and J.W.M. designed and executed the research studies. C.E. assisted with *Streptococcus pyogenes* strain construction and reagents. J.B.D. assisted with experimental design. N.C.K. and M.J.S. assisted with plasmid construction and *Streptococcus pyogenes* strain construction. R.E.W. and J.W.M. wrote the manuscript.

## METHODS

### RESOURCE AVAILIBILITY

#### Lead Contact

For further information and requests for resources and reagents, please contact Joshua Modell, Department of Molecular Biology and Genetics, Johns Hopkins School of Medicine, Baltimore, MD.

#### Materials Availability

All materials generated for this study are available upon request and without restrictions from the Lead Contact, Joshua Modell.

### EXPERIMENTAL MODEL AND SUBJECT DETAILS

#### Microbes

*Staphylococcus aureus* cells were grown at 37°C, unless otherwise indicated, in Bacto Brain-Heart infusion (BHI) broth with shaking at 220 RPM. During outgrowths from stationary phase preceding phage treatments, BHI was supplemented with calcium chloride at 5 mM to allow phage adsorption and with 1 mM IPTG to allow expression from inducible promoter Psparc2 when necessary. Antibiotics were used at the following concentrations for strain construction and plasmid maintenance in *S. aureus*: tetracycline, 5 µg/mL; chloramphenicol, 10 µg/mL; erythromycin, 10 µg/mL; spectinomycin, 250 µg/mL.

*Streptococcus pyogenes* cells were grown at 37°C, unless otherwise indicated, in Bacto Todd-Hewitt broth with 2% yeast (ThY) without shaking. During outgrowths from stationary phase preceding phage treatments, ThY media was supplemented with calcium chloride at 5 mM and 2 mg/ml sodium bicarbonate. During outgrowths from stationary phase preceding phage top agar treatments, dialyzed ThY media (preparation detailed below) was supplemented with calcium chloride at 5 mM and 2 mg/ml sodium bicarbonate. Antibiotics were used at the following concentrations for strain construction and plasmid maintenance in *S. pyogenes:* chloramphenicol 3 µg/mL, spectinomycin 100 µg/mL, kanamycin 150 µg/mL.

#### Phages

*Staphylococcus aureus* phage ɸNM4g4 was amplified on RN4220 and stored in BHI at 4°C. *Streptococcus pyogenes* phages A1 and AP1.1 were amplified on strain C13, a derivative of SF370 prophage-cured strain CEM1ΔΦ^33^ and a kind gift of Andrew Varble, which has a spontaneous deletion of spacers 1-5 in its CRISPR array, including deletion of the natural spacers targeting phage A1 and ϕAP1.1. Amplified Streptococcal phage stocks are stored in ThY at 4°C.

### METHOD DETAILS

#### Plasmid construction

See Supplemental Materials for strains, plasmids, cloning notes, and oligos used in this study.

#### Gibson assembly

Gibson assemblies were performed as described^42^. Briefly, 100 ng of the largest dsDNA fragment to be assembled was combined with equimolar volumes of the smaller fragment(s) and brought to 5 µL total in dH2O on ice. Samples were added to 15 µL of Gibson Assembly master mix, mixed by pipetting and incubated at 50°C for 1 hour.

For electroporation into Staphylococcus aureus RN4220, samples were drop dialyzed in dH2O for 30 minutes to 1 hour, and 5 µL were electroporated into 50 µL electrocompetent RN4220 *S. aureus* cells. For heat shock transformation into Escherichia coli, 5 µl of Gibson assembly was added to chemically competent Dh5a, cells, incubated for 2 hours at 37°C with shaking, then plated on the appropriate antibiotic.

#### Streptococcus pyogenes Todd-Hewitt dialysate media preparation

To optimize phage infection assays in *S. pyogenes*, we utilized a media dialysis recipe from Zabriskie 1964. We mixed 100 ml water, 30g Bacto Todd Hewitt Broth media, and 20g Bacto yeast extract in a 500 ml Erlenmeyer flask, then microwaved and agitated until homogenous and clear (2-5 min). This media was placed into 11 inches of dialysis tubing (Spectra Por S/P 3; molecular weight cut-off 3.5 kDa), sealed, and floated in 2 L distilled water for 1 hour. The water was replaced and the media dialyzed for another 2 hours, then the tubing was transferred to a 4 L beaker filled with distilled water and dialyzed overnight in the cold room for 16 hours. The dialyzed media was transferred to a 1L beaker, brought up to 1L volume with distilled water, and autoclaved for 45 minutes. Soft top agar for phage infection assays was made by adding 0.2% agarose to Thy-D and autoclaved for 45 minutes.

#### Transformation in Streptococcus pyogenes

*S. pyogenes* cells were made electrocompetent through outgrowth of an overnight culture diluted 1:20 to an OD of ∼0.3 in 50 ml, followed by centrifugation at 4000 x g at 4° C for 20 minutes. The supernatant was decanted and the cell pellet was washed with 50 mL cold 10% glycerol, and pelleted at 4000 x g at 4° C for 15 minutes. The cell pellet was resuspended in 2 mL cold 10% glycerol, split between two 1.5 mL Epppendorf tubes, then centrifuged at 6000xg at room temperature for 1 minute. The cells were washed two additional times with 1 mL cold 10% glycerol, with centrifugation at 6000 x g for 1 minute. The washed cell pellet was resuspended in 500 µl 10% glycerol, then separated into 50 µl aliquots. 5 µl of >100 ng/µl plasmid DNA was mixed with the electrocompetent cells, then pipetted into a pre-chilled 0.1cm cuvette (Biorad, 165-2089). The cuvette was dried with a Kimwipe, then pulsed at 2.5kV/cm, 200 ohms, and 25 µF. 950 µl of pre-warmed BHI was immediately added to the cuvette, then moved to a 1.5 mL Eppendorf and incubated at 37°C without shaking for 3 hours. After incubation, the cells were struck out onto BHI plates with the appropriate antibiotic.

#### Markerless allelic exchange

To create mutations in the native *Streptococcus pyogenes* host chromosome, as well as within lysogenized phage AP1.1, we performed markerless allelic exchange. Each allelic exchange construct was designed with 500-1000bp left and right homology arms amplified from the wildtype background, flanking the mutation to be made, and cloned via Gibson assembly into the vector pCRK/pCRS, generous gifts of Dr. Kevin McIver. Each plasmid was first transformed into *E. coli* Dh5a, then isolated from 1-10 mL overnight culture using the Qiagen miniprep kit (Thermo Fisher Scientific) according to manufacturer’s instructions. The purified plasmid was transformed into *Streptococcus pyogenes* strain SF370 using the electroporation protocol described below and plated onto BHI plates with the appropriate antibiotic and incubated at 30°C. After transformation, colonies were inoculated into 1 mL liquid media with antibiotic and incubated at the restrictive temperature of 37°C. The resulting overnight culture was plated onto BHI plates with the appropriate antibiotic and incubated at the restrictive temperature. Single cross-overs were confirmed by PCR and inoculated into 1 mL BHI with antibiotic and grown overnight at 37°C without shaking. These stocks were then frozen down in 10% DMSO and re-struck onto BHI plates with antibiotic and incubated at 30°C. Three single colonies were picked and passaged at 30°C in 10 mL liquid media without antibiotic overnight, then 10 µl of this culture was transferred to 10 mL of BHI broth without antibiotic overnight. This passaging step was repeated, and the resultant population was plated onto BHI plates without antibiotic. To assess double cross-over status and excision of the plasmid, 100 individual colonies from each plate were patched onto both plates without antibiotic and plates with. Colonies which were sensitive to the antibiotic but still grew on plain BHI were lysed in 100 µl 1X PBS and PlyC (1 µg/ml final concentration), then checked for successful mutant generation using PCR.

#### Phage A1 mutant construction

Primers and template used to construct phage mutants are detailed in Supplementary Data 1. Each A1 allelic exchange construct was designed with 500-1000bp left and right homology arms amplified from the wildtype background, flanking the mutation to be made, and cloned via Gibson assembly into the vector pC194. This construct was transformed into *S. pyogenes* without a targeting spacer (JW3886) to avoid CRISPR targeting. Additionally, a selection plasmid with *crRNA* leader-repeat-spacer-repeat sequence on pLZ12 backbone was created, with the *crRNA* spacer targeting wild-type, unrecombined phage A1. This plasmid was then transformed into *S. pyogenes Δtracr-L* without targeting spacer (JW3917). To generate A1 recombinants, the S. pyogenes strain containing recombination plasmid was diluted to OD=0.1 in 1 ml Thy supplemented with 5 mM calcium chloride and 2 mg/ml sodium bicarbonate. After a ∼2 hour incubation at 37°C without shaking, and at an OD between 0.3-0.4, the culture was infected with wildtype phage A1 at MOI=5. The infection was incubated at 37C without shaking until culture clearance, which took ∼2-4 hours. Cell debris was pelleted at 8000 x g for 1 minute and supernatant was filtered through a 0.2µm filter. The phage was stored at 4C until recombinant selection.

To select for recombinants, the *S. pyogenes* strain containing a phage A1 WT targeting spacer, and optionally, a control strain with just a pLZ12 control plasmid, was outgrown by diluting an overnight culture 1:10 in 1 mL Thy-D supplemented with 5 mM calcium chloride and 2 mg/ml sodium bicarbonate, and letting grow at 37°C to an OD of 0.5-1. 500 µl of cells were mixed with 5 ml 0.2% Thy-D (or Thy) top agar supplemented with 5 mM calcium chloride and 2 mg/ml sodium bicarbonate, pipetted with a serological to mix and 1.5 ml of this mix was plated on BHI bottom agar with the appropriate antibiotic. To validate phage A1 recombinants, 4 plaques per plaqued plate were selected and pipetted into 20 µl Thy. 2 µl of the resuspended phage was added to 10 µl phage lysis buffer (1 µl 20 mg/ml Proteinase K into 100 µl 1X PBS), incubated at 55°C for 20 minutes and 98°C for 10 minutes. The phage lysate was amplified with primers of interest using primers outside the homology arms and validated with Sanger sequencing.

#### S. aureus miniprep protocol

1-1.5 mL of an overnight culture, unless otherwise indicated, was pelleted and resuspended in 250 µL Qiagen Buffer P1. 10-20 µL Lysostaphin (Ambi Products LLC, LSPN-50, 100 µg/mL final) was added and the cells were incubated without shaking at 37°C for ∼20 minutes. Following lysis, plasmids were isolated using the QIAGEN Spin Miniprep kit according to the manufacturers protocol, beginning with addition of P2. DNA was eluted from each column in 30 µL RNAse-free water.

#### Liquid growth interference assay

Overnight cultures of *S. aureus* were diluted to OD = 0.025 in BHI broth supplemented with 5 mM calcium chloride and antibiotics and grown for 1 hour and 15 minutes shaking at 37°C. Cultures were normalized to the OD=0.1 and phage ɸNM4g4 was added to the appropriate MOI. After inverting to mix, 150 µL of each culture were added to a flat-bottom 96-well plate (Grenier 655180), and the plate was incubated at 37°C with shaking in a TECAN Infinite F Nano+ with OD600 measurements recorded every 10 minutes for 24 hours.

For *Streptococcus pyogenes* assays, overnight cultures of *S. pyogenes* were diluted back 1:10 in fresh Thy media supplemented with 5 mM calcium chloride and 2 mg/ml sodium bicarbonate and grown for 1 hour and 30 minutes without shaking at 37°C. Cultures were normalized to the OD=0.05 and phage A1 (and derivatives) was added to the appropriate MOI. After inverting to mix, 200 µL of each culture was added to a flat-bottom 96-well plate (Grenier 655180), and the plate was incubated at 37°C without shaking in a TECAN Infinite F Nano+ with OD600 measurements recorded every 10 minutes for 24 hours.

#### Top agar interference assay

For *Staphylococcus aureus* assays, 100 µL of *S. aureus* overnight cultures were added to a 50 mL Falcon tube. 6 mL of BHI top agar (0.75% agar) supplemented with calcium chloride (5mM final concentration) and IPTG (1mM final) was added to each tube. After swirling to mix, the cells and top agar were poured onto a BHI 1.5% agar plate and rocked gently to create a bacterial lawn. The plate was incubated for 15 - 30 minutes at RT. 3.5 µL of 8 10-fold serial dilutions of phage ɸNM4g4 in BHI broth were spotted on top of the bacterial lawn using a multichannel pipette. After a 30 minute incubation at room temperature, the plates were moved to a 37°C incubator overnight.

For *Streptococcus pyogenes* assays, overnight cultures of *S. pyogenes* were diluted back 1:10 in fresh Thy-D media supplemented with 5 mM calcium chloride and 2 mg/ml sodium bicarbonate. After a 2-4 hour outgrowth, 200 µl of cells were mixed with 1.8 ml Thy-D top agar (Thy-D with 0.2% agarose), 5 mM calcium chloride and 2 mg/ml sodium bicarbonate. This mixture was inverted to mix, and 1.5 ml was pipetted onto BHI plates. The plate was incubated for 15 - 30 minutes at RT to dry the top agar. Phage A1 or AP1.1 was diluted 10-fold 8 times, and 3.5 µL was spotted on top of the bacterial lawn using a multichannel pipette. After a 30 minute incubation at room temperature, the plates were moved to a 37°C incubator overnight.

#### Promoter activity fluorescence assays

To quantify promoter activity of Pcas, the promoter was fused to GFP and expressed on a plasmid also containing *tracr-L*. To assay fluorescence, 200 µL of each overnight culture, with cells in late stationary phase, was spun down in a 1.5 mL Eppendorf tube at 6,000 rpm for one minute. Cell pellets were resuspended in 1 mL of 1X PBS and 150 µL was transferred into a clear, flat-bottomed 96-well plate (Grenier 655180). Measurements for absorbance (at 600 nm) and GFP fluorescence were recorded.

#### PCR conditions

PCR was performed with Phusion HF DNA polymerase using 5X Phusion Green Reaction Buffer (Thermo). Each reaction contained 10 µL buffer, 4 µL dNTPs, 0.5 µL each of 100 µM forward and reverse primers, 10-50 ng template, 0.5 µL polymerase and nuclease-free water to 50 µL. Three-step cycling was performed under the following conditions: 98°C for 30 seconds, 34 cycles of [98°C 5 seconds, 45-72°C 15 seconds, 72°C for 30s/kb], 72°C 10 minutes, hold at 10°C.

#### RNA extraction

To extract *S. aureus* RNA for Northern blot analysis, 7.5E8 cells were spun down, resuspended in 150 µL 1X PBS (10X stock, Corning, 46-013-CM) and lysostaphin (60 µg/ml final concentration), and incubated at 37°C for 5 minutes. Cells were processed from overnight cultures unless otherwise specified. To the whole cell lysate, 450 µl Trizol (Zymo, R2071) and 600 µL 200 proof ethanol were added, samples were vortexed and RNA was extracted using the Direct-Zol Miniprep Plus spin column according to the manufacturer’s protocol (Zymo, R2071). Samples were eluted in 50 µL Ambion RNAse-free water (ThermoFisher, AM9937).

To extract *S. pyogenes* RNA for Northern blot analysis, 7.5E8 cells from an overnight culture were spun down, resuspended in 150 µl 1X PBS and PlyC (1 µg/ml final concentration), and incubated at room temperature for 10 minutes. After lysis, the RNA extraction protocol was followed as detailed above.

#### Radioactive Northern Blot

Total RNA (3-10 µg) was mixed 1:1 with 2X Novex sample buffer (Thermo, LC6876), boiled at 94°C for 3 minutes and placed on ice for 3-5 minutes. Samples were loaded onto a 15% TBE-Urea gel (MINI-Protean, Bio-rad, 4566053) and run at 150 V for 3 hours. A Hybond N+ membrane (GE lifesciences, 45000854) and 6 sheets of 3 mm Whatman cellulose paper (Sigma Aldrich, WHA3030861) were pre-soaked for 5 minutes in room temperature 0.5X TBE, then assembled into a sandwich of: 3 layers Whatman paper, Hybond membrane, TBE-Urea gel, and 3 more layers of Whatman paper. Blotting was performed using a Trans-blot Turbo (Bio-Rad) at 200 mA for 30 minutes. The membrane was then pre-hybridized in 10 mL ExpressHyb (Clontech, NC9747391) at 44°C in a rotating oven for 1 hour, and probed in fresh hybridization buffer overnight at 44°C, with rotation, using probes labeled with P32. The membrane was washed once with 2X SSC/0.1% SDS, and once with 1X SSC/0.1% SDS each for 10 minutes at RT. The gel was then removed from its casing, wrapped in plastic wrap and exposed to a phosphor screen overnight. The phosphor screen was scanned and imaged using the Typhoon FLA9500 (GE Healthcare). 4.5S RNA was used as a loading control (stability of 4.5S across genetic backgrounds was verified by qPCR). For Northern blots performed using *S. pyogenes* cells, 5S RNA was used as a loading control. Sequences for oligos used to probe *crRNA*, *tracrRNA* and *4.5S* RNA (oJW2313, oJW1991, oJW2172) are listed in Supplementary Materials.

#### Western blot

1.8E8 *S. aureus* cells were resuspended in 1X PBS supplemented with lysostaphin (60 µg/ml final concentration) and incubated at 37°C for 20 minutes. Cells were processed from overnight cultures unless otherwise specified. Whole cell lysate was mixed 1:1 with 2x Laemmli solution (Bio-rad, 1610737) supplemented with B-mercaptoethanol (55 mM stock, ThermoFisher, 21985023) at a final concentration of 1.3 mM, and boiled at 98°C for 10 minutes. Samples were loaded onto a 4-20% Tris-glycine gel (MINI-Protean TGX Pre-cast, Bio-rad, 4561095) and run at 200 V for 15 min-1 hour. A PVDF membrane (Bio-rad) was hydrated with methanol for 15-30 seconds and pre-wet alongside a stack of blotting paper for 3-5 minutes in 1X Transfer buffer. A blotting sandwich was assembled consisting of six layers (one stack) of filter paper, the PVDF membrane, Tris-glycine gel, and another stack of filter paper. Samples were transferred onto a nitrocellulose membrane with the Trans-blot turbo (Bio-Rad, 1704150), with high molecular weight transfer settings (1.6 A for 10 minutes), and the membrane was stained with Ponceau for 5 minutes to perform total protein normalization. The membrane was blocked with 5% nonfat dry milk in TBST for 1 hour, then probed with a 1:1000 dilution of Cas9 monoclonal antibody (Cell Signaling, 7A9-3A3) for 2 hours at RT, or at 4°C overnight. The membrane was washed with 1X TBST buffer 3x for 10 minutes at RT, then probed a 1:10,000 dilution of HRP (Pierce, PA174421) secondary antibody for 1 hour at room temperature. The membrane was washed as before, then visualized on the Odyssey Fc (LICOR).

To analyze *S. pyogenes* Cas9 levels, ∼1.8E8 cells were resuspended in 1X PBS supplemented with PlyC (1 µg/ml final concentration) and incubated at room temperature for 10 minutes. After lysis, the Western protocol is followed as detailed above.

#### qPCR

Purified total DNA was diluted to 0.5-1 ng/µL and 5 ng was used as input in an 8 µL volume. 1 µL of 10 µM forward/reverse primers were used, and 10 µL of 2X PowerUp SYBR mastermix (ThermoFisher, A25742). qPCR was performed with cycling conditions: 50°C for 2 minutes, 95°C for 2 minutes, 39 cycles of [60°C 1 minute], followed by a melt curve: 65°C to 95°C, incrementing 0.5°C every 5 seconds. qPCR primer sequences are provided in Supplementary Materials.

#### Burst curve assay

To determine latency period and infection cycle timing for phage A1, we used a burst curve assay. Overnight cultures of *S. pyogenes* were diluted back 1:10 in Thy media supplemented with 5 mM calcium chloride and 2 mg/ml sodium bicarbonate and grown for 1 hour and 30 minutes without shaking at 37°C. Cultures were normalized to OD=0.1 in a total volume of 10 mL and phage A1 was added at an MOI of 1. The culture was inverted 10X to mix, then aliquoted into 10 1 ml eppendorfs and incubated at 37°C without shaking. One tube was removed at each timepoint and then centrifuged at 6000 xg for 1 minute. 10 µL of PFU supernatant was removed, serially diluted, and plaqued on a top agar lawn of (non-interfering) *Streptococcus pyogenes* cells.

#### Infective centers assay

To determine percentage of cells surviving the first infection cycle, we used an infective centers assay. Overnight cultures of *S. pyogenes* were diluted back 1:10 in fresh Thy media supplemented with 5 mM calcium chloride and 2 mg/ml sodium bicarbonate and grown for 1 hour and 30 minutes without shaking at 37°C. Cultures were normalized to OD=0.05 in a total volume of 1 mL and phage A1 (and derivatives) was added at an MOI of 0.1. Cultures were incubated at 37°C without shaking for 10 minutes to allow for phage adsorption and injection, then centrifuged at 6000 xg for 1 minute. The supernatant was removed and the remaining cell pellet was resuspended in 1 mL 1X PBS. The centrifugation and resuspension steps were repeated, and then the appropriate volume of cell resuspension (obtained by prior experiments determining which volume resulted in between 50-500 countable plaques) was added to 150 µl of C13 (non-interfering) *Streptococcus pyogenes* cells. Quickly, 1.4 mL of dialyzed Todd-Hewitt soft agar with 5 mM calcium chloride and 2 mg/ml sodium bicarbonate was added to the mixture and plated on BHI bottom agar. Effective centers of infection (ECOI) was then calculated by dividing the number of infective centers per ul formed from infection of the targeting strain by the number of infective centers per ul formed from infection of the non-targeting strain and then subtracting this from 1. This value then represents the percentage of cells surviving the first round of infection.

#### One-step growth curve (OSGC) assay

To estimate phage latency period and burst size, we used a one-step growth curve (OSGC) assay. Overnight cultures of *S. pyogenes* were diluted back 1:10 in Thy media supplemented with 5 mM calcium chloride and 2 mg/ml sodium bicarbonate and grown for 1 hour and 30 minutes without shaking at 37°C. Cultures were normalized to OD=0.05 in a total volume of 500 µL and phage A1 (and derivatives) was added at an MOI of 3. Cultures were incubated at 37°C without shaking for 10 minutes to allow for phage adsorption and injection, then centrifuged at 6000 xg for 1 minute. Cells were resuspended in 500 µl Thy and 2 mg/ml sodium bicarbonate, then diluted 1:1,000 in the same media. After dilution, 10 µl was set aside to plate for infective centers before the remainder of the culture was centrifuged at 10,000 xg for 1 minute. 20 µl of the supernatant was then set aside for plaque quantification. The appropriate volume of either cell suspension or PFU supernatant (obtained by prior experiments determining which volume resulted in between 20-200 countable plaques) was added to 150 µl of C13 (non-interfering) *Streptococcus pyogenes* cells. Quickly, 1.4 mL of dialyzed Todd-Hewitt soft agar with 5 mM calcium chloride and 2 mg/ml sodium bicarbonate was added to the mixture and plated on BHI bottom agar.

#### Induction of lysogens

To isolate AP1.1 phages, overnight cultures of the appropriate lysogen were diluted 1:10 into 1 ml Thy and grown at 37°C to an OD∼0.3 (∼2 hours). Cells were induced with 0.5 µg/ml mitomycin C for 4 hours, then cells were centrifuged at 6000 xg for 1 minute and supernatant was filtered through a 0.20 µm sterile syringe filter (Corning, 431229).

#### Lysogeny assay

To estimate percentage of cells undergoing lysogenic conversion after ɸAP1.1 infection, overnight cultures of *S. pyogenes* were diluted back 1:10 in fresh Thy media supplemented with 5 mM calcium chloride and 2 mg/ml sodium bicarbonate and grown for 1 hour and 30 minutes without shaking at 37°C. Cultures were normalized to OD=0.2 in a total volume of 180 µL and ∼4000 PFUs in 20 µl phage AP1.1 (and derivatives) were added. Cultures were incubated at 37°C without shaking for 60 minutes to allow for phage adsorption and injection, then the full volume (200 µl) was plated on BHI + 50 µg/ml spectinomycin using glass beads.

### QUANTIFICATION AND STATISTICAL ANALYSIS

#### Promoter activity fluorescence assays

Measurements for absorbance (at 600 nm) and fluorescence (excitation wavelength = 485 nm; emission wavelength = 535 nm) were taken using a TECAN Infinite F Nano+. For each experimental strain, promoter activity was measured as (Fe)/(Ae) - (Fc)/(Ac) where F = fluorescence, A = absorbance, e = experimental strain and c = non-fluorescent control strain.

#### Area under the curve analysis

Measurements for bacterial growth during Acr-phage infection were taken using absorbance at 600 nm with a TECAN Infinite F Nano+ every 10 minutes for 20 hours. The generated curves were quantified using area under the curve analysis, with baseline set to Y=0. Analysis was conducted using Prism 7.

### KEY RESOURCES TABLE

*Separate file

## Supplemental Figures

**S1.**
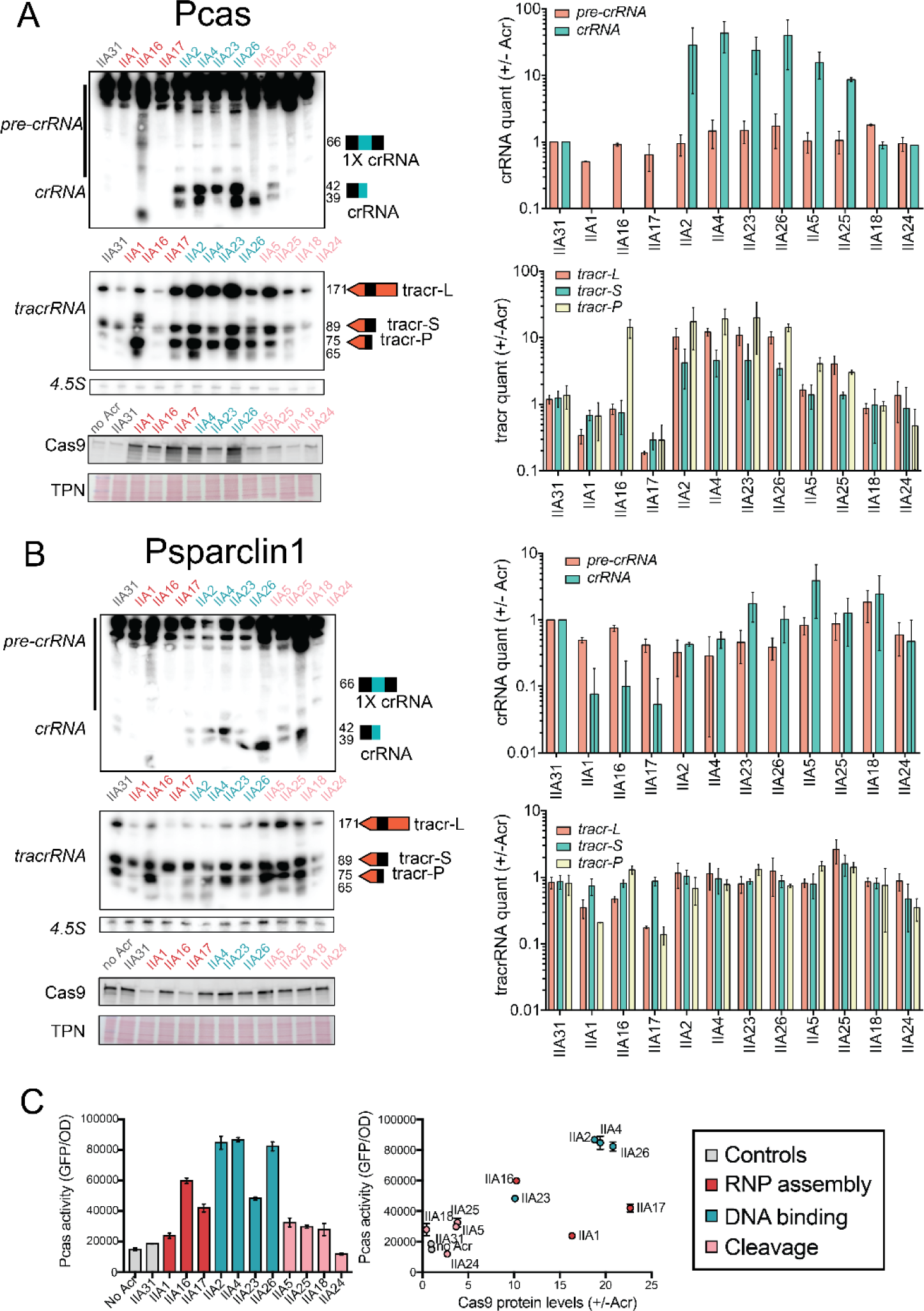
Anti-CRISPRs induce Cas expression through disruption of *tracr-L* repressor. Pcas (A) and Psparclin1 (B) Northern blots for *crRNAs* and *tracrRNAs* and Western blots for Cas9 in *S. aureus* cells treated with 1 mM IPTG for 120 minutes to induce Acr expression. Graphs show Northern blot quantifications in biological duplicate or triplicate. C) GFP fluorescence normalized to OD600 was measured in cells expressing a Pcas-GFP transcriptional reporter following a 120 minute 1 mM IPTG treatment. Raw values in biological triplicate are plotted on the left panel, and a scatterplot of Cas9 protein levels in Pcas cells from Fig. 1D vs Pcas-GFP levels for the indicated Acrs are shown in the right panel.

**S2.**
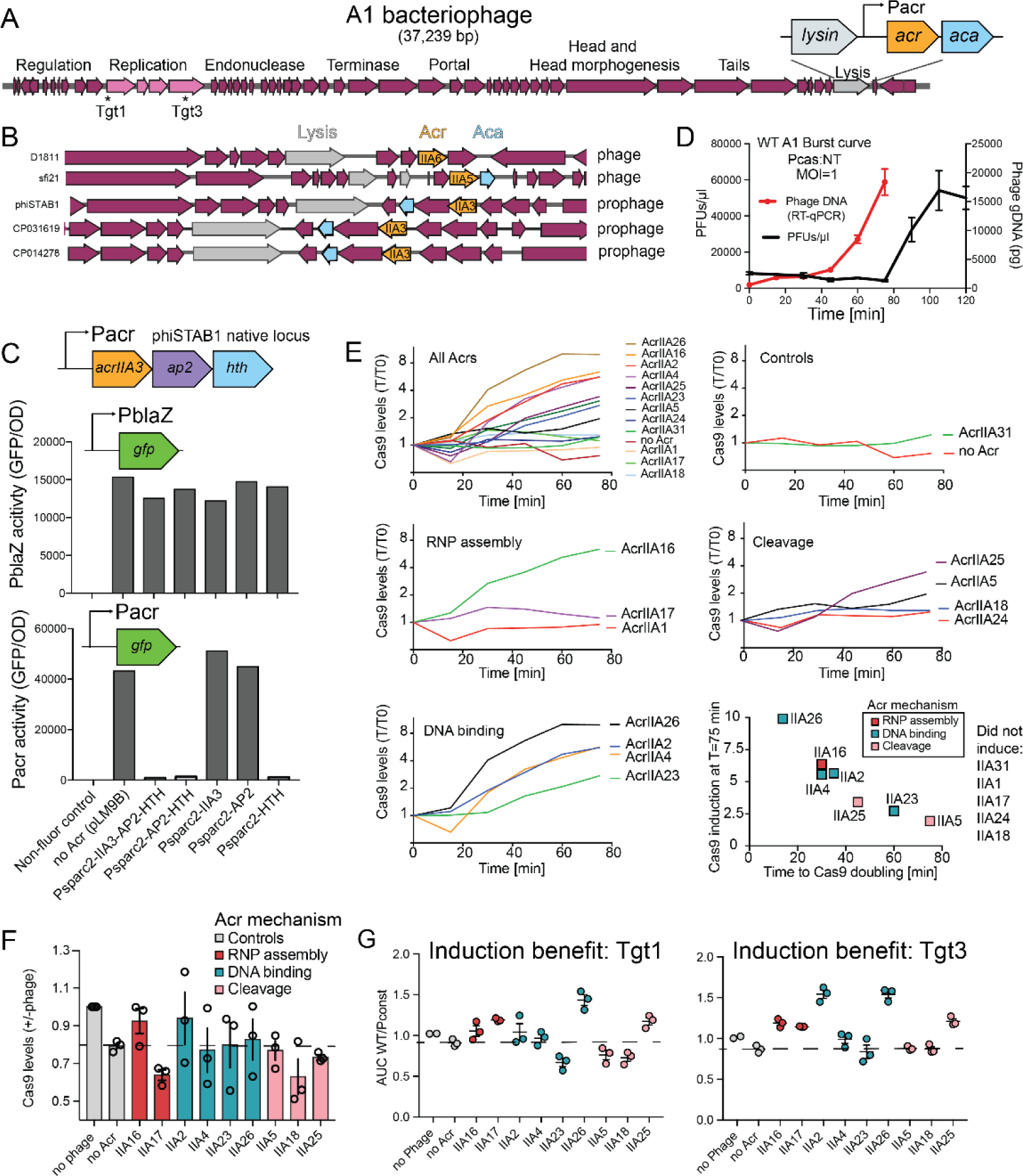
Anti-CRISPRs expressed from bacteriophage A1 induce Cas expression during an active phage infection. A) Phage A1 genome, labeled with the locations of targeting sites for spacers Tgt1 and Tgt3 and the insertion of the Acr-aca operon. B) *Streptococcal* phages often host Acrs downstream of the lysis module; the locations of lysins (grey), Acrs (yellow), and Acas (blue) are shown for each genome. C) Reporter assay validating phiSTAB1 *hth* as an anti-CRISPR associated (Aca) gene. Fluorescence normalized to OD600 was calculated in cells expressing the indicated phiSTAB1 genes from one plasmid and either PblaZ-GFP (control promoter) or Pacr-GFP from a second plasmid. D) A burst-curve assay was performed on non-interfering Pcas cells (C13) treated with phage A1 at MOI=1. Phage DNA replication was monitored by qPCR (red line) and infectious particles by a PFU assay (black line) at the indicated time points. E) Western blot quantifications for Pcas cells with a single spacer targeting A1 infected at MOI=2 for with the indicated A1-Acr phages. Each 15 minute timepoint was normalized to T=0. Cas induction at 75 minutes was plotted against the time required for Cas9 levels double (bottom right panel). F) Cas9 levels were measured by Western blot in Pconst cells infected with the indicated A1-Acr phage at MOI=2 for 75 minutes, normalized to a no phage control. Dotted line represents Pconst-Cas9 levels of cells infected with WT phage A1. G) The induction benefits for the indicated A1-Acr phages was calculated as in Fig. 2E for cells with single A1-targeting spacers (Tgt1, Tgt3). MOIs of infections provided in Supplemental Data 1.

**S3.**
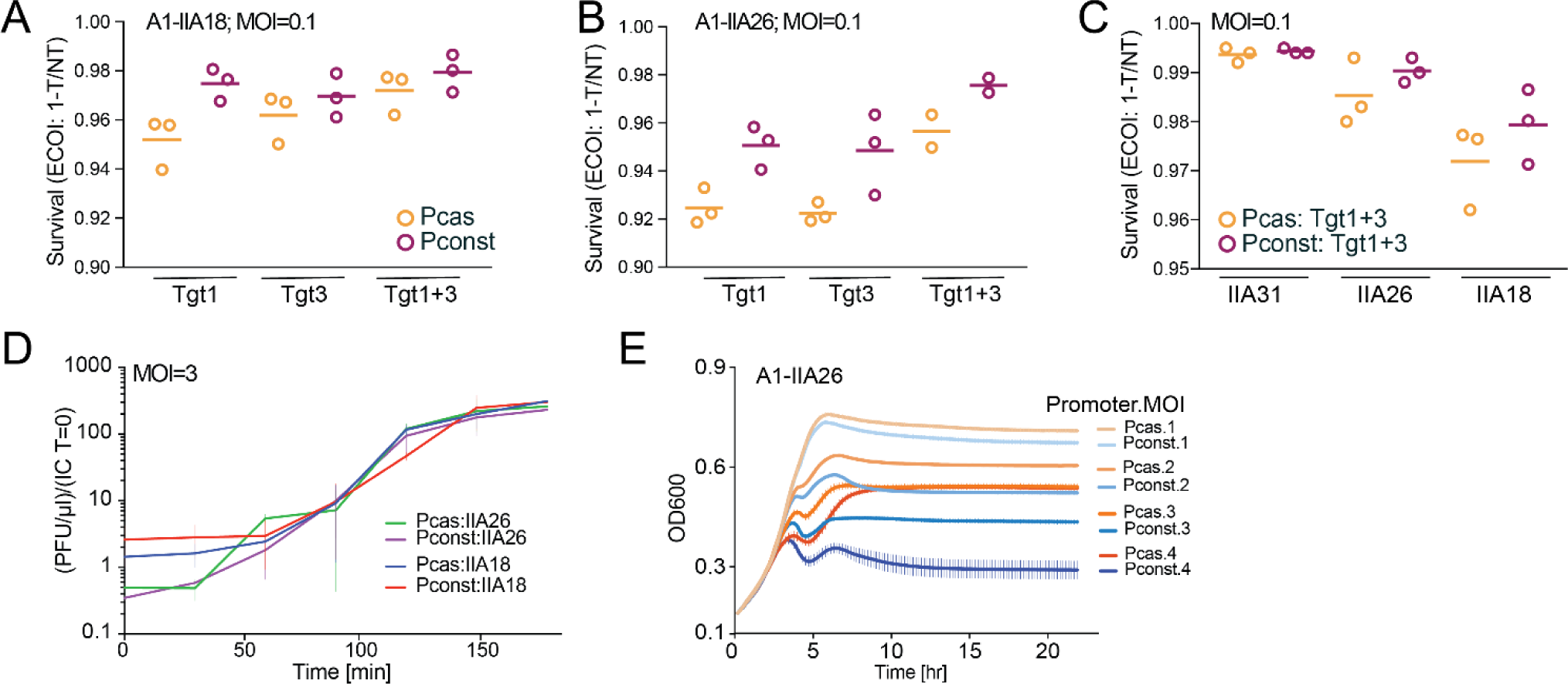
The benefit of CRISPR-Cas induction is MOI dependent and exacerbated by multiple infections. A-C) Effective centers of infection (ECOI) assay at MOI=0.1 for Pcas or Pconst cells with the indicated phages and spacers. The proportion of infected cells that survive the primary infection is plotted on y-axis (1-(targeting PFUs/non-targeting PFUs)). D) OSGC assay for Pcas or Pconst interfering cells (Tgt1+3) infected at MOI=3 withphage A1-IIA26 or A1-IIA18. PFUs were normalized by infective centers to determine the burst size of each infection. E) 22-hour growth curves for Pcas and Pconst cells (Tgt1+3) infected with MOIs 1- 4.

**S4.**
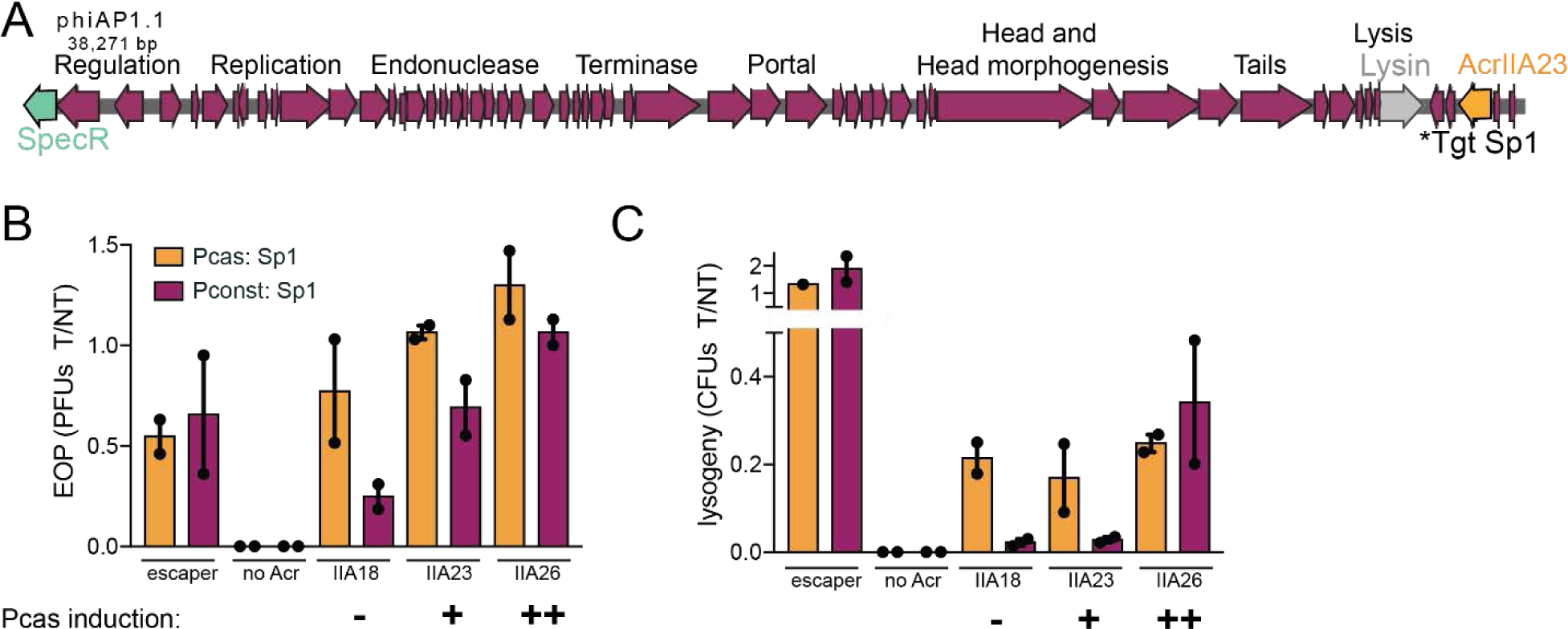
PhiAP1.1 lysis and lysogeny is impacted by CRISPR-Cas induction. A) The phiAP1.1 genome is shown including the positions of the spectinomycin resistance gene^22^ (green), lysin (grey), AcrIIA23 (orange), and targeting site for Sp1. B) Top agar interference assay for phage AP1.1 (“IIA23”) or AP1.1 variants expressing Acr-IIA18 or -IIA26 plated on Pcas or Pconst cells with a spacer (Sp1) targeting AP1.1. C) A lysogeny assay was conducted on the same strains and phages as in (A). Lysogeny rates were calculated as the ratio of lysogens formed in targeting (T) divided by non-targeting (NT) cells.

